# Enhanced Multiplexed Single-Cell RNA-Sequencing for Accurate Detection of Treatment Effects Without Batch Correction in the Avian Embryonic Model

**DOI:** 10.1101/2025.09.30.679584

**Authors:** Stephanie Maupetit-Mehouas, Felipe Maurelia Gaete, Pouria Hosseinna, Nicolas Allègre, Yoan Renaud, Jonas Cruzel, Claire Chazaud, Charlène Guillot

## Abstract

Single-cell RNA sequencing has revolutionized our ability to explore cellular heterogeneity and is a powerful tool to study the impact of environmental perturbations on multiple cell states. However, environmental perturbations can be subtle, and the associated biological effects could be masked by experimental noise and bioinformatic processing, especially when the samples are generated separately. Multiplexing strategies have been developed to label each sample and process them together to reduce experimental noise, but existing multiplexing methods often fall short for non-human and non-mouse models in established single-cell RNA sequencing protocols like the BD Rhapsody technology. To address this gap, we combined Lipid Modified Oligonucleotide (LMO) cell tagging with the BD Rhapsody platform to achieve efficient and scalable multiplexing of early embryonic chick cells under different environmental conditions. This species-agnostic LMO approach overcomes limitations of antibody-based multiplexing methods that are often restricted to human and mouse systems, and the only multiplexing option available with the BD Rhapsody system. In our study, we successfully compartmentalized and multiplexed chick embryonic cells under different treatment conditions, analyzing up to 40,000 viable cells per experiment. This strategy minimized experimental noise, eliminating the need for bioinformatics-based batch correction. As a result, we were able to achieve high-quality transcriptomic profiling with minimal loss of critical biological information and identified subtle biological differences that were masked when using data integration pipelines. Our workflow provides an adaptable, robust solution for LMO-tagged single-cell analyses of complex non-human models with the BD Rhapsody technology and opens new avenues for developmental biology research by accurately capturing treatment-induced effects in embryonic tissues.

## 1. Introduction

Single-cell RNA sequencing (scRNA-seq) has emerged as a powerful tool for exploring biological heterogeneity in human and model organisms at the single-cell level (Cheng et al., 2023; Perkel, 2021). Recent advances in scRNA-seq have enable the incorporation of cells from multiple samples into single libraries through multiplexing strategies, thereby improving sample throughput while minimizing technical batch effects, library preparation time, and overall costs (Mylka et al., 2022; Xie et al., 2024). Such strategies allow for direct comparison of different samples under genetic or environmental perturbations while minimizing bioinformatic processing (McFaline-Figueroa et al., 2024; Srivatsan et al., 2020). Sample multiplexing is generally achieved by exploiting either preexisting genetic diversity or by introducing sample-specific barcodes before pooling, a process commonly referred to as “hashing.” Several approaches have been developed for this purpose, including oligo-labeled antibodies, chemical oligo-tagging and lipid-modified Oligonucleotides (LMOs) (Gehring et al., 2020; McGinnis et al., 2019; Stoeckius et al., 2018). Nevertheless, most of these strategies have been optimized and validated primarily in human and mouse systems, limiting their direct applicability to non-traditional or embryonic model organisms. Among them, LMOs have emerged as a versatile tool for sample multiplexing, with potential compatibility with most single-cell RNA sequencing technologies (Brown et al., 2024; McGinnis et al., 2019). However, LMO-based cell labelling introduces additional processing steps prior to single-cell encapsulation, directly affecting cell viability and quality of the analysis (Brown et al., 2024). This impact is particularly relevant for samples with low cell numbers or high complexity, such as *in vivo* embryonic animal model tissues. To this date, LMOs have not been used to label highly heterogeneous early chick embryonic cells.

The BD Rhapsody single-cell RNA sequencing platform offers a notable advantage for processing samples with high cell heterogeneity and RNA content, enabling the generation of robust, high-quality data from *in vivo* animal tissues (Ulbrich et al., 2023). Compared with other platforms such as 10x Genomics Chromium, BD Rhapsody demonstrated superior performance with small-sized cells, making it particularly suitable for developmental context where cell dimensions are reduced (Colino-Sanguino et al., 2024a). Currently, the most widely used multiplexing strategy for this platform relies on antibody-based cell tagging with species-specific pan-cell antibodies, allowing the analysis of up to 40000 cells per experiment (Ulbrich et al., 2023). Nevertheless, this approach faces important limitations: antibody-based tagging depends on epitope recognition, which may fail to capture the full cellular diversity within the samples (Salcher et al., 2024). Such constrains increase the risk of missed cell populations, especially considering that available reagents are currently restricted to human and mouse samples. Notably, BD Rhapsody has not been applied to multiplexing strategies in the chick model for scRNA-seq.

By leveraging LMO-based multiplexing, which can target all animal species and cell types, we can overcome the limitations associated with antibody-dependent strategies and perform multiplexing of embryonic samples under different environmental conditions. In addition to enabling the analysis of up to 40,000 cells per experiment, this microwell-based scRNA-seq method produces high-quality data from cell suspensions with a minimal starting point of 50% viable cells. This is achieved through a post-loading washing step that removes dead cells and environmental RNA from the media (Ulbrich et al., 2023). Based on these advantages, we combined the BD Rhapsody scRNA-seq method with LMO tagging to perform multiplexed analysis of early chick embryonic cells exposed to different treatments, a context characterized by dissociation-sensitive heterogeneous cell populations. While a similar approach was recently used to perform single-cell sequencing capture of tumor cells in mice (Colino-Sanguino et al., 2024a), no such experimental procedure has been performed yet for the early chick embryonic cells under different environmental conditions.

The chick model is an exquisite model for developmental and biomedical research (Asai et al., 2021; Garcia et al., 2021). Used in the clinics for teratoma potency of small molecules, it is considered a suitable model for the analysis of cancer cells to diverse treatments/chemicals (Corsini et al., 2021; Javed et al., 2023). Thanks to its easy accessibility, it has been a great model to study the role of environmental perturbations during development (Bień et al., 2025; Darmawan et al., 2025; de Cristo Soares Alves et al., 2023; Ibrahim et al., 2025) and allows to easily access and study early developmental programs such as gastrulation and body axis formation (Guillot et al., 2021; Mongera et al., 2019). It is in this framework that we wanted to analyze the role of homocysteine treatment on body axis formation. Indeed, previous work showed that adding homocysteine to the avian embryo, to mimic folate deficiency in humans, induces neural tube-like defects, indicating an impairment of the neural tube morphogenesis process. However, how it impacts the associated developmental program and cell states during embryogenesis is not fully understood. Here, we decided to combine targeted homocysteine treatment with multiplexed single-cell RNA sequencing to identify the impact of homocysteine treatment on the transcriptomic cell state of the cells involved in posterior body axis formation.

In our study, we combine the benefits of pan-cell tagging from LMOs and the analysis of only viable cells to achieve high-quality single-cell RNA sequencing of vertebrates’ embryonic samples under varying treatment conditions simultaneously, with a capacity of up to 40,000 living cells per experiment.

## Results

### Uniform labeling of embryonic chick cells with Lipid Modified Oligonucleotides

Lipid Modified Oligonucleotides (LMOs) offer a versatile sample barcoding solution compatible with various cell types or nuclei (McGinnis et al., 2019; Mylka et al., 2022). Its hydrophobic extremity can insert into accessible plasma membranes, facilitating pooled single-cell sample processing from varying treatment conditions. Comprising a Lignoceric amide-modified anchor DNA oligo and a palmitic amide-modified co-anchor DNA oligo, these lipid-modified oligos seamlessly embed into cell or nucleus membranes, providing a landing pad for DNA barcodes with complementary 5’ sequences in its ‘handle’ region (McGinnis et al., 2019).

To assess LMO insertion into membranes of chick cells, we designed a fluorescent probe capable of hybridizing to the ‘handle region’ at the 3’ terminal end of the Anchor, specifically marked with Atto fluorescent label 550 (Figure 1A Right). Insertion efficiency was analyzed via Flow cytometry (BD Melody). We tested three LMO final concentrations (200nM, 20nM, and 2nM) to label dissociated cells (Supplementary Table 1). At the fabricant-recommended LMO concentration of 200nM, we observe better labeling of cells, shown by the uniform distribution on the histogram (Figure 1B). A lower concentration reveals a biphasic behavior on the histogram, with highly and less labeled cells (Figure 1B). Using the fabricant-recommended LMO concentration of 200nM, we achieved LMO-labelling of 79% of living single cells with 15% of mortality (Figure 1C). Using confocal imaging, we confirmed that we can detect single cells alive (Hoechst positive) and labeled with LMO (Atto 550 positive) after the dissociation and insertion (Figure 1D). Our findings demonstrate that LMOs can efficiently label viable embryonic chick cells. Subsequently, we investigated whether this method enables sample multiplexing for the BD Rhapsody single-cell RNA sequencing.

**Figure 1:**
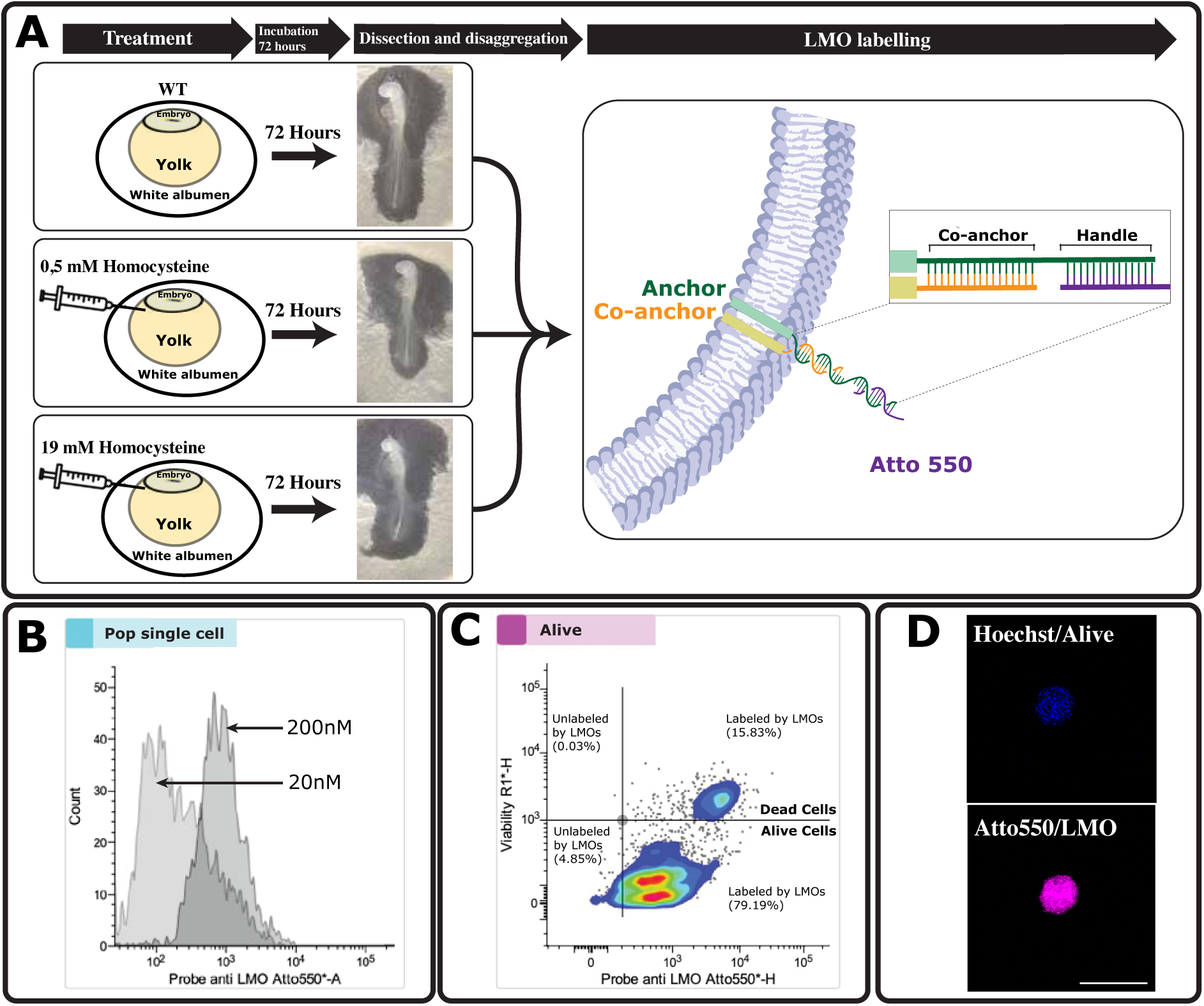
Labeling of embryonic chick cells with Lipid Modified Oligonucleotides (LMO). **A**. (Left) Schematic representation of the experimental procedure showing chicken embryos treated with homocysteine at incubation time and incubated for 72hours. Schematic representation of dissociated Cell labeled with LMO using anchor and co-anchor probes together with a Handle fused to the Atto550 dye for fluorescence detection. **B**. Histogram of single-cell population labeled with LMOs at 20nM and 200nM, analyzed by flow cytometry. **C**. Density plot showing the distribution of LMO-labeled cells within live and dead populations, based on Live/Dead488 and Atto 550 intensities in single cells measured on a BD FACS Melody^TM^. Approximately 15% are dead (upper right), 79% are alive and LMO-labeled (lower right), and 4.8% are alive but non-labeled (lower left). **D**. Representative confocal image of a dissociated live cell (Hoechst positive) labeled with LMOs (Atto550 positive), acquired at 20X magnification on a Zeiss LSM 980.

### Performing LMO Multiplexing Single-Cell RNA Sequencing with BD Rhapsody

For labeling each cell from a specific sample, a unique Sample Barcode (SBC) was synthesized with unique 8-nucleotide barcodes. These sequences feature a 5’ region complementary to the ‘handle’ of the anchor, followed by validated barcode sequences(McGinnis et al., 2019) and a 3’ poly A tail to attach to the rhapsody bead (refer to Supplementary Table 2).

To perform LMO labeling of chick embryonic cells, cells are dissociated and labeled per sample by a specific mixture of Anchor-coAnchor and SBC (See star method). After incubation on ice, excess of LMOs were removed by centrifugation steps. After assessing final cell viability after LMO labeling, 13000 viable cells of each sample are loaded onto the cartridge. The steps of cell capture were performed following those outlined in the BD Rhapsody protocol and the library preparation steps (Figure 2).

**Figure 2:**
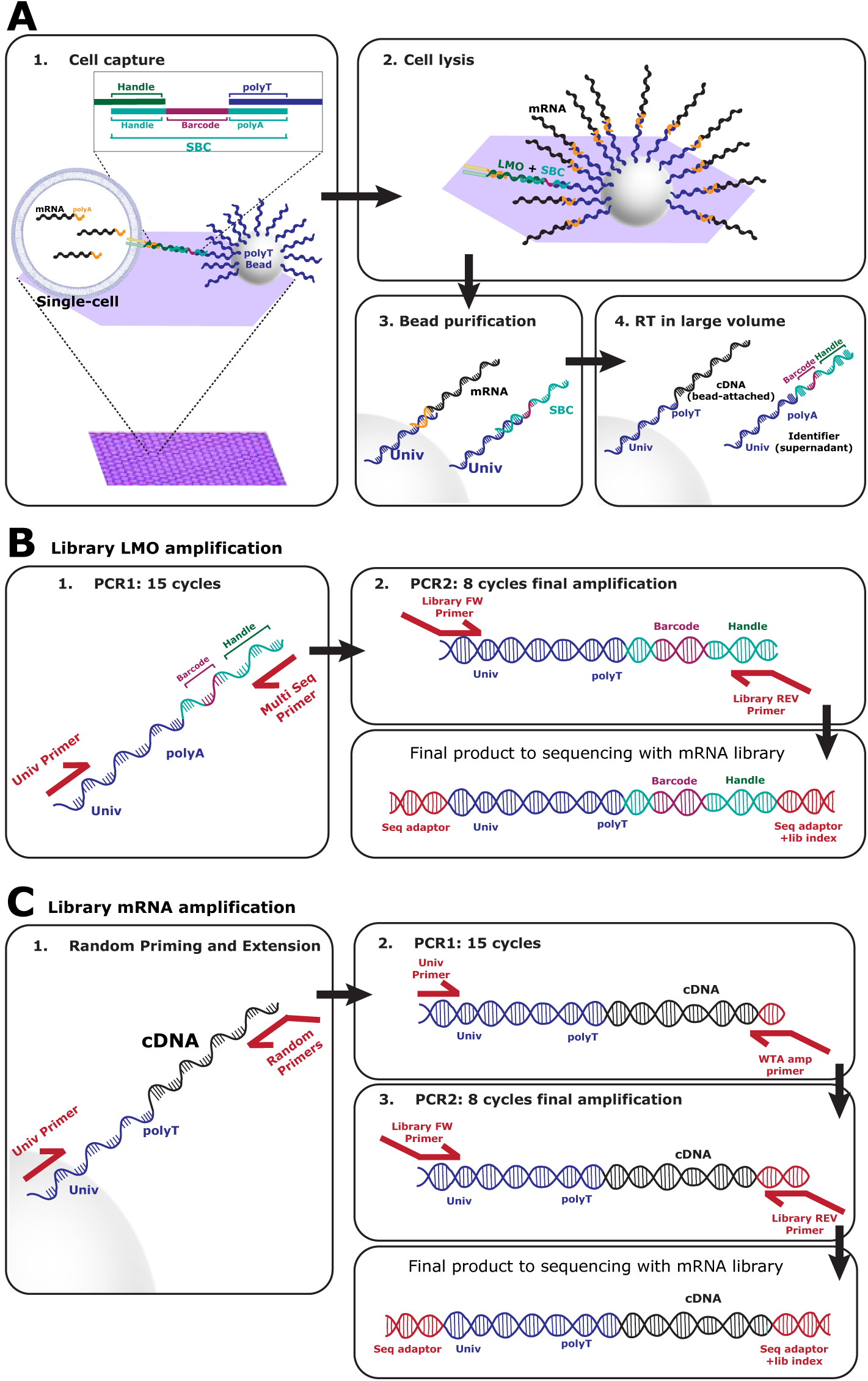
Steps of mRNA and LMO capture from single-cell-derived embryonic chick tissue using the Rhapsody Cartridge. A. Diagram illustrating the main steps of cell capture in the BD Rhapsody system. Unlike standard capture methods, the sample barcode construct (SBC) contains a poly-A region, enabling the identifier to hybridize to the beads after cell lysis. This feature ensures its retention during purification and amplification during the reverse transcription step, thereby allowing efficient capture and downstream analysis. **B**. LMO library amplification by PCR on the supernatant after lysis, enabling the amplification and sequencing of barcode regions for cell identify assignment. Custom primers are listed in Supplementary Table 3.**C**. mRNA library amplification by PCR on the purified beads, required for whole-transcriptome library preparation as indicated in the BD Rhapsody protocol.

### Adapting the library preparation from the BD™ AbSeq protocol to retrieve the LMOs

mRNAs and LMOs from individual cells were captured on Rhapsody beads through a specific sequence at their extremities, which includes a polyT tail (Figure 2A). This design enables the identification of the mRNA population for a specific cell and, with the help of LMOs, associates the data with a specific biological sample.

After capturing mRNAs and LMOs on beads, reverse transcription and exonuclease treatments are performed following the established BD protocol. The supernatant from the beads is preserved and processed separately to generate a library specific to the LMOs, allowing for the barcoding of cells (Figure 2B and Star method). Here, we used a specific primer named Multi Seq Primer that is complementary to the Handle region to perform the PCR1 step which also contains the complementary region to allow the binding of Primer PCR2 (Supplementary table 3). Simultaneously, a transcriptome library is generated using beads loaded with cDNA (Figure 2C). This process involves amplification by random priming and extension, followed by two PCR steps. To sequence the two libraries, we pair-end 150 bp sequencing on a high-through Illumina platform. Depth of sequencing for the LMO library was 2000 reads/cells and 50 000 reads/cells for Whole Transcriptome Analysis library.

### Reads alignment and quality controls

BD Rhapsody provides an online platform i.e. “Seven Bridges” with in-house analysis pipeline, allowing users to check the sequencing quality, align and annotate the reads, and generate the necessary output files required for downstream analysis. We adapted the VDJ function alignment to retrieve the LMO sequences within the BD Rhapsody™ WTA Analysis Pipeline (Figure 3A). To do so, we provided the FASTQ files from the WTA and LMO sequencing, the reference genome, the transcriptome annotation, and the Barcode sequences as a supplemental reference file. Output files include expression matrices filtered or unfiltered, BAM files, and a pipeline report indicating the quality of the sequencing and alignment (supplementary HTML file).

**Figure 3:**
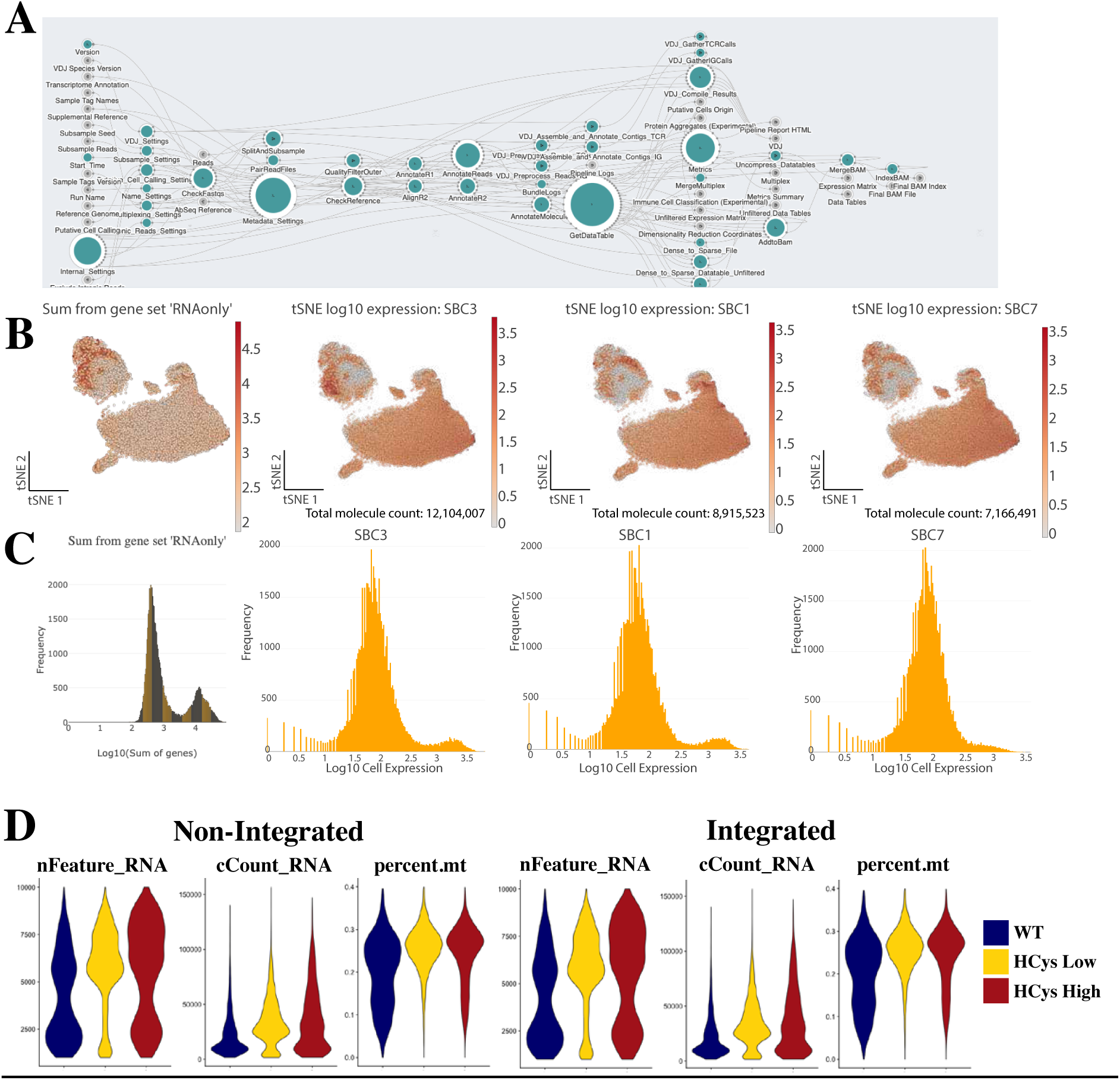
Quality control of sequencing and read alignments. **A**. Schematic representation of the modules used in the BD Rhapsody™ WTA Analysis Pipeline on the Seven Bridges platform. Modules used for aligning mRNA and LMO reads to the chick genome and identifying known sample barcodes are shown in green. Additional complementary modules available for optional analysis are depicted in grey. **B**. tSNE plots illustrating the distribution of cells across the three analyzed samples. The left plot shows the aggregate gene expression retrieved from the WTA library for all samples. The right plots depict the expression levels of individual sample barcodes retrieved in the analysis. **C**. Histograms representing the frequency distribution of gene expression (left) and sample barcode expression (right) per cell. **D**. Quality control of the barcoded cells retained for the analysis depicting the expression levels of the number of genes per cells nFeatures_RNA), the number of reads per cells (nCount_RNA) and the percentage of mitochondrial genes expressed per cells (percent.mito) in each individual sample barcodes. WT: Wild-type, HCys Low: 0,5mM Homocysteine-treated, HCys High: 19mM Homocysteine-treated.

Quality controls of the alignment indicate successful amplification of both mRNA and LMOs. Specifically, 96.41% of reads passed the quality filters, with 75.6% of cellular reads aligning uniquely for mRNA, while 97.16% of reads passed quality filters, and 79.87% of cellular reads aligned uniquely for LMOs (Figure 3B, C, and supplementary HTML file).

The t-SNE plots (Figure 3B, left) depict the sum of gene sets from the WTA library for all three samples combined with 28,146 total genes detected. Good expression levels of each sample’s barcodes were also retrieved for each sample (Figure 3B, right).

Notably, the expression of the sample barcodes follows a normal distribution, with most cells expressing around 100 barcodes (log-transformed values of 2, Figure 3C). This distribution confirms efficient barcoding and successful retrieval of barcodes together with expected mRNA retrieval (Figure 3C, left).

### Seurat analysis of the LMO-tagged multiplexed cells

To analyze the data, we utilized the published Seurat pipelines to demultiplex the data, assign the correct LMO identity, and perform quality control (Colino-Sanguino et al., 2024b). Applying stringent filters for LMO numbers per cell (>100 barcodes per cell), we retrieved 16,385 single cells barcoded for analysis (Figure 3D). High-quality cells were defined as those with less than 0.4% mitochondrial gene content and expression of 1,000 to 10,000 genes. After filtering, we retained 11,982 high-quality single-barcoded cells across the three samples for downstream analysis (Figure 3D).

### Cell multiplexing enables the detection of subtle transcriptomic shifts under environmental perturbations

Environmental perturbations often elicit subtle transcriptomic changes that could be overlooked in conventional analysis. In standard scRNAseq experiments, samples are processed separately, which introduces technical variations that manifest as batch effects and reduced comparability across conditions. Bioinformatic integration pipelines are typically employed to mitigate these effects. However, such an approach can also obscure subtle transcriptomic changes after treatment (McFaline-Figueroa et al., 2024; Srivatsan et al., 2020). To overcome this limitation, we conducted a multiplexing strategy that minimizes batch effect and experimental variability, enabling direct comparison of samples without requiring integration steps. To test whether integration compromises the detection of subtle changes, we compared datasets from WT and homocysteine-treated embryos before and after applying the SCT transform integration algorithm within Seurat. In the non-integrated analysis, UMAPs revealed many overlapping regions, but also treatment-specific regions (Figure 4B, arrow). In contrast, after SCT-based integration, these distinct regions were no longer observed (Figure 4B right), suggesting that some biological effects were masked by computational integration. To further validate this observation, we performed a complementary analysis of cluster composition (Figure 4C). For each cluster, we calculated the proportion of cells originating from each sample and tested whether these proportions significantly deviated from the uniform distribution expected under multiplexed loading. In the non-integrated data, the treatment shows a change in the sample composition across multiple clusters —including Paraxial Mesoderm, Lateral Mesoderm, NMPs, Blood, and Neural Crest Progenitors— whereas after SCT-based integration, only the Blood cluster remained significantly shifted. These results indicate that integration can attenuate treatment-linked compositional signals that multiplexed acquisition reveals. Altogether, these findings indicate that minimizing variability at the data generation stage through multiplexed single-cell capture and sequencing enhances the sensitivity to detect fine-grained transcriptional differences between treatments, which might otherwise be masked by computational batch correction.

**Figure 4:**
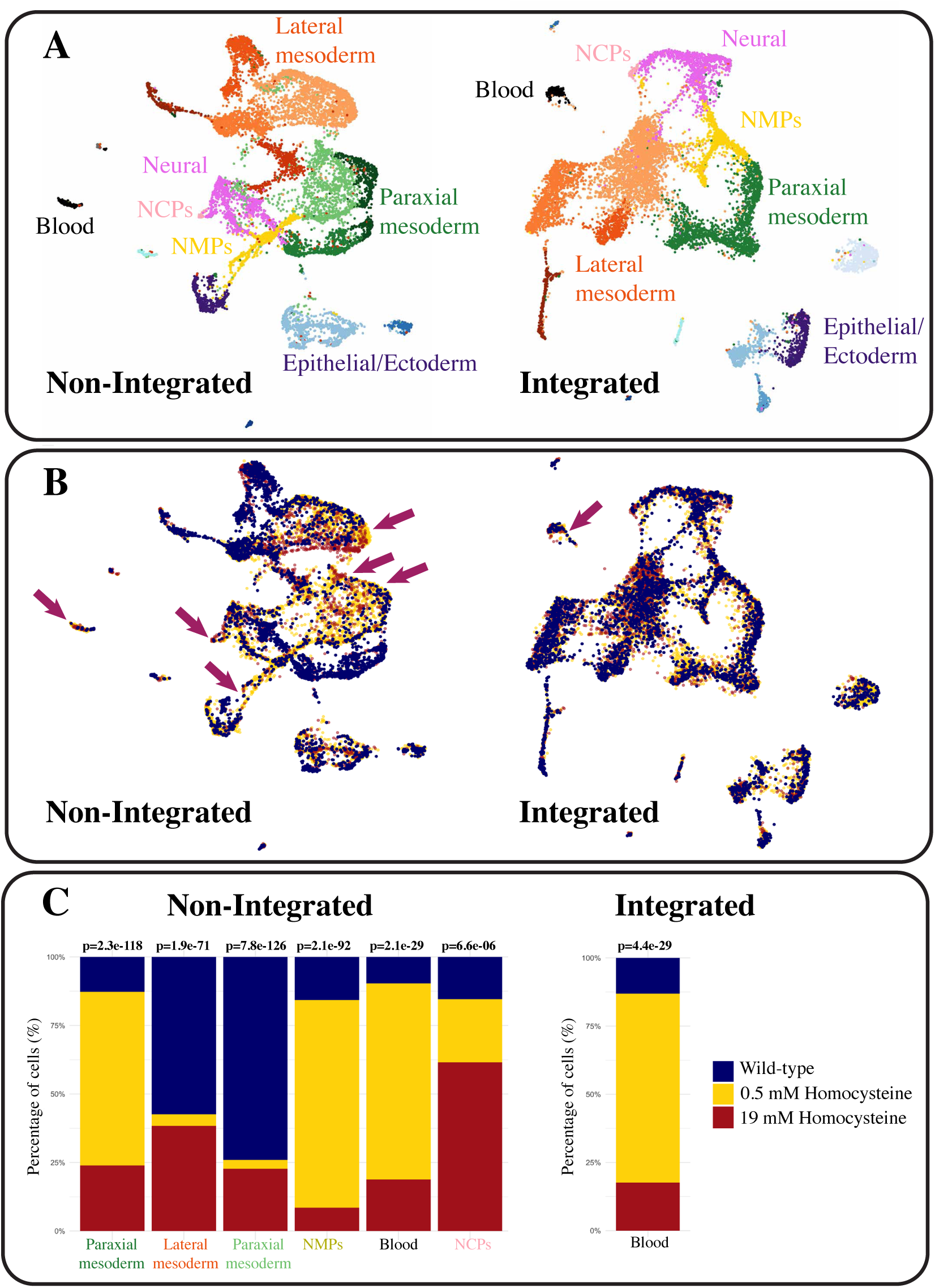
Non-integrated method reveals discrete single-cell differences in gene expression. **A**. UMAP plots showing cell populations identified by specific markers in clustered groups analyzed without integration and with integration using the SCT transform Seurat function. **B**. UMAP plots displaying the distribution of cells from WT and treated embryos (yellow and red). Arrows indicate regions where distinct differences are observed. C. Comparison of cluster composition showing differences in the percentage of cells identified after treatment in both methods.

Overall, we successfully combined for the first time BD Rhapsody Technology with LMO labelling of chick embryonic cells to perform multiplexed single-cell RNA sequencing. This approach enabled us to analyze transcriptomic state changes in ∼16,000 high-quality cells contributing to posterior body axis formation. Our findings demonstrate that this strategy is particularly well-suited to detect subtle biological changes, which are crucial for understanding developmental errors induced by environmental perturbations.

## Discussion

Single-cell RNA sequencing (scRNA-seq) has revolutionized the study of cellular heterogeneity, providing unparalleled insights into complex biological systems (Perkel, 2021). However, different multiplexing strategies have a profound impact on data quality, scalability, and applicability across species. To date, most validated protocols have relied on antibody-based tagging, such as TotalSeq HashTag antibodies, which are restricted to human and mouse systems (ref). In contrast, our study demonstrates that Lipid-Modified Oligonucleotides (LMOs) constitute a versatile and specie-agnostic labelling method that can be easily combined with the BD Rhapsody microwell-based platform. This combination enables robust multiplexing of chick embryonic cells during early development, thereby addressing critical challenges in assessing the impact of environmental perturbations on dissociation-sensitive and heterogeneous embryonic tissues.

Compared to antibody-based approaches, LMO labelling offers several advantages. First, its independence from epitope recognition avoids potential biases coming from limited antibody availability or uneven cell surface marker exposition, thereby offering coverage for the highly heterogeneous early embryonic tissue. Second, the combination of LMO labelling with BD Rhapsody’s washing steps ensures that only viable cells can contribute to downstream analysis, effectively minimizing background signal from ambient RNA. At the recommended concentration of 200nM of LMO label, we achieved uniform labelling of ∼80% of viable cells, while maintaining scalability to up to 40.000 cells per cartridge. By retaining only cells with a high barcode count (>100), we obtained 16,000 cells for downstream analysis. Our sequencing depths— 2,000 reads per cell for LMO libraries and 50,000 reads per cell for transcriptome libraries—meet or exceed industry benchmarks for multiplexed workflows (Xie et al., 2024). These results are consistent with their prior use on PBMCs and cell lines demonstrating that LMO labelling achieves a high signal-to-noise ratio when optimized (McGinnis 2019; Brown 2024). Importantly, all the cell states retrieved in the LMO multiplexed combined with BD Rhapsody were previously found when using no multiplexing combined with indrop single cell sequencing, albeit with deeper sequencing depth, and more genes were recovered with this novel technology. (Guillot et al., 2024, 2021). LMOs have also been successfully applied in other vertebrate models, including zebrafish brain development (Brown et al., 2023), supporting their broad applicability across species. In the avian model, Gill et al. (2024) recently applied MULTI-seq to study intestinal development at later embryonic stages, although without specifying the use of LMO or CMO anchors (Gill et al., 2024). Our work extends these findings by not only establishing a validated workflow for multiplexed scRNA-seq in early chick embryos—a critical window for body axis formation with highly heterogeneous populations that had not been addressed previously—but also by demonstrating, for the first time, the integration of LMO barcoding with the BD Rhapsody platform. This novel combination provides a robust and scalable approach for dissociation-sensitive embryonic tissues, expanding the applicability of both LMOs and BD Rhapsody beyond human and mouse systems.

A central challenge of scRNA-seq is distinguishing true biological variation from technical noise. When comparing different data types, batch correction algorithms are often used to mitigate technical artifacts (Büttner et al., 2019; Hu et al., 2025).

However, these algorithms can sometimes overcorrect, removing genuine biological information. Overcorrection can make downstream analyses problematic and often result in incorrect conclusions. In our study, LMO barcoding allowed us to compartmentalize cells from the outset, thereby reducing technical variability and eliminating the need for computational batch correction. This approach ensures precise sample identification and comprehensive transcriptomic profiling. Indeed, we observe in our analysis that all cells occupy similar spaces in the UMAP plot without any batch effect corrections. More importantly, when using batch effect correction algorithms, some cell types were merged and subtle variations after treatment were hidden. This suggests that our multiplexing strategy provides a significant advantage, preserving subtle transcriptomic changes -such as those induced by environmental treatments-that might otherwise be corrected by computational integration, thereby enabling more accurate biological interpretations.

From a biological perspective, our results provide new evidence on how homocysteine treatment perturbs early embryonic development in chicken embryos. Previous studies have associated elevated homocysteine with neural tube defects (Kobus-Bianchini et al., 2017; Taparia et al., 2007; Wenstrom et al., 2000), impaired neural crest delamination and migration (Brauer and Rosenquist, 2002; Melo et al., 2017), and altered vascular function (Boot et al., 2004; Bourckhardt et al., 2019; Latacha and Rosenquist, 2005; Oosterbaan et al., 2012). In line with this, we identified several cell populations linked to these processes, underscoring the broader impact of homocysteine treatment on early embryonic lineages. Importantly, both integrated and non-integrated analyses consistently revealed a population of cells associated with blood progenitors -one of the best-documented lineages affected by homocysteine treatment during embryo development-further supporting its relevance as a teratogenic molecule. Further, it is interesting to note that it is only in the non-integrated analysis that we identify a role of homocysteine in the neural crest and posterior body axis cell types, validating this approach to recover more subtle effects of environmental perturbations on developmental programs. Nevertheless, the precise molecular mechanism by which homocysteine exerts these effects remains speculative, and further work is needed to dissect the direct effects of this molecule and its indirect consequences in disrupting the one-carbon metabolism during early development.

Together, our findings demonstrate that integrating LMO tagging with the BD Rhapsody platform provides a powerful approach for high-quality scRNA-seq of complex samples including early heterogeneous embryonic tissues. This methodology offers significant potential for expanding single-cell studies to include non-mouse or human organisms and broadens the scope of multiplexing workflows to challenging dissociation-sensitive systems and environmentally challenged conditions where changes may be more subtle than genetic perturbations.

## Conclusion

We present a reliable and adaptable workflow for multiplexed scRNA-seq of embryonic chick cells using LMO tagging integrated with the BD Rhapsody platform. This method enables high-quality analysis of dissociation-sensitive, heterogeneous tissues, scales to 40,000 cells, and is broadly compatible with non-human models. By minimizing batch effects during capture, library preparation, and sequencing, it eliminates the need for bioinformatic batch correction and preserves subtle biological differences found in environmentally challenged conditions. This approach offers a robust tool for accurate transcriptomic comparisons in developmental biology and expands single-cell research to previously underexplored systems.

## Supporting information

STAR methods

## Resource availabily

Requests for further information and resources should be directed to and will be fulfilled by the lead contact, Charlene Guillot.

## Materials availability

This study did not generate new unique reagents.

## Data and code availability

The single-cell RNA sequencing datasets generated and analysed during the current study have been deposited in the NCBI Gene Expression Omnibus (GEO) repository under accession number GSE303866. This paper does not report original code. Any additional information required for re-analysing the datasets reported in this paper is available from the corresponding authors upon request.

## Limitations of Study

A first limitation of our study lies in the reproducibility of the injection procedure, as each egg presents intrinsic differences that may introduce variability on the results after treatment. To mitigate this, we mixed multiple samples to minimise the impact of inter-embryo variation. A second limitation concerns the number of cells analyzed: although the dataset is robust, it can be bias by the capacity of the BD Rhapsody cartridge on capture different cell populations equally. Nonetheless, the identification of expected cell populations is consistent with our previous dataset generated without multiplexing and with a drop-based single cell platform (Guillot et al., 2021), supporting the accuracy of the current data. Finally, in terms of biological interpretation, this work represents an initial step. While our analyses reveal transcriptomic differences between treated and untreated embryos, further validation—through hybridization chain reaction (HCR) or by functional assays—will be necessary to confirm the observed changes and to better understand the mechanisms underlying homocysteine’s effects during early embryonic development.

## Ethical statement

This study did not require ethical approval, as it involved only early-stage *Gallus gallus* embryos collected before hatching, in compliance with European Directive 2010/63/EU.

## Acknowledgements

The authors thank Dr. Krzysztof Jagla from the Institute of Genetics and Reproduction for their helpful discussions, suggestions, and comments. We are also grateful to the BD team and the Seven Bridges support members for their assistance and technical help. The authors also would like to acknowledge the outstanding technical services provided by the Institute of Genetics, Reproduction and Development platforms: SC3 – Single Cell & Cell Culture Facilities, the Bioinformatics Platform (BIM), and the Clermont Confocal Imaging Platform (CLIC). We also thank the administrative and technical shared support staff of the institute for their constant assistance.

## Credit authorship contribution statement

**Stephanie Maupetit-Mehouas**: Writing – review & editing, Writing – original draft, Visualization, Methodology, Investigation. **Felipe Maurelia Gaete**: Writing – review & editing, Visualization, Software, Investigation, Formal analysis, Data curation. **Pouria Hosseinna:** Writing – review & editing, Software, Formal analysis, Data curation, **Nicolas Allegre**: Methodology, Investigation. **Yoan Renaud**: Software, Formal analysis, Data curation. **Jonas Cruzel**: Methodology, Investigation. **Claire Chazaud:** review & editing, Supervision, Resources. **Charlene Guillot**: Writing – review & editing, Writing – original draft, Project administration, Supervision, Resources, Methodology, Investigation, Formal analysis, Data curation, Funding acquisition, Conceptualization.

