## Supplementary material for "Enhanced Multiplexed Single-Cell RNA-Sequencing for Accurate Detection of Treatment Effects Without Batch Correction in the Avian Embryonic Model": STAR methods

### EXPERIMENTAL MODEL AND STUDY PARTICIPANT DETAILS

### METHOD DETAILS

**Experimental model, treatments, and sample preparation**

Fertilized *Gallus gallus* eggs (JA57 strain) were obtained from EARL Les Bruyères (Dangers, France) and incubated under standard conditions. To induce experimental treatments, embryos received a supplementation of homocysteine consisting of 20 µL of a 19 mM solution (high dose) or 20 µL of a 0.5 mM solution (low dose), injected into the subembryonic region of the yolk, while control embryos were injected with 20 µL of Hank’s Balanced Salt Solution (HBSS, Sigma-Aldrich, cat#H8264). At 72 hours post-incubation (Hamburger–Hamilton stages 18–19), embryonic tissues from treated and control embryos were collected in parallel under identical conditions to ensure comparability. For each experimental condition, two anatomical regions were dissected from four embryos: the tail bud (two embryos), representing the posterior-most structures, and the posterior end (two embryos), which included the last formed somite, presomitic mesoderm, posterior neural tube, and tail bud. Tissues were briefly rinsed in cold HBSS and enzymatically dissociated in Accutase (Gibco, Cat#A1110501) for 15 min at 37 °C, followed by gentle mechanical trituration. The resulting suspension was filtered through a 40 µm cell strainer, and cell density and viability were evaluated using a hemocytometer. Finally, cells were centrifuged at 350 × g for 5 min at 4 °C and resuspended in 0.25% BSA in PBS.

**Validation of LMOs insertion into the membrane of dissociated chicken cells by flow cytometer**

For LMO labelling, 5 × 10⁵ dissociated cells were incubated on ice with Anchor LMOs for 5 min, followed by the addition of an equal concentration of co-Anchor LMOs for another 5 min. Three concentrations of Anchor/co-Anchor LMOs were tested in parallel: 200 nM, 20 nM, and 2 nM. To simultaneously monitor cell viability and LMO incorporation, cells were co-incubated with Fixable Viability Stain 660 (FVS660, BD Biosciences; 1:200 from stock) and a complementary Atto550-labelled probe (0.1 µM). After incubation, cells were centrifuged at 500 × g for 10 min at 4 °C, resuspended in HBSS, and analyzed on a BD FACS Melody™ cytometer.

Controls included unstained cells, FVS660-only, probe-only, and LMOs + probe, which were used to define gating strategy and compensation. Data were acquired and processed using standard analysis workflows (see Figure 1).

**Single cell experiments using LMOs and BD Rhapsody**

Single-cell dissociation protocols were optimized to achieve >90% viability and minimize doublets before capture. For LMO labelling, 2.5 × 10⁵ cells from each tissue were incubated in PBS supplemented with 200 nM Anchor and 200 nM tissue-specific Sample Barcode (SBC) oligonucleotide, each SBC being dedicated to one tissue type. After agitation and 5 min on ice, 200 nM co-Anchor was added for an additional 5 min. Cells were washed twice by centrifugation at 500 × g for 10 min at 4 °C. Based on viability counts, 16,000 live cells per sample (up to 40,000 total) were loaded into a primed BD Rhapsody™ cartridge (BD Biosciences) according to the manufacturer’s protocol. Cell capture beads were dispensed into each microwell, and cells were lysed to release mRNA and SBC-tagged LMOs, which were subsequently captured by the beads. After bead washing, reverse transcription was performed, followed by exonuclease I treatment to remove unbound primers.

**Library preparation and sequencing**

- **LMO barcode library**

To recover SBC-derived barcode information, reverse transcription products not bound to the beads were collected from the supernatant. A first PCR amplification was carried out using the MULTI-seq additive primer, which specifically recognizes the SBC handle sequence. The expected amplicon size (~138 bp) was verified after AMPure XP bead purification (Beckman Coulter, Cat#A63880) using a TapeStation 4200 (Agilent). A second PCR was then performed to add Illumina P5 and P7 adapters, using custom reverse primers (Supplementary Table 3). The resulting final library had a theoretical size of ~158 bp.

- **Whole transcriptome library (WTA)**

Beads containing captured mRNA were processed with the BD Rhapsody™ WTA Amplification Kit according to the manufacturer’s instructions without modifications. This workflow generated cDNA libraries representative of whole-transcriptome profiles for each captured cell.

**Sequencing**

LMO and WTA libraries were pooled at equimolar concentrations and sequenced on an Illumina NovaSeq platform (paired-end 150 bp, Novogene). The sequencing depth was configured to achieve ~1,000 reads per cell for the LMO barcode library and ~50,000 reads per cell for the WTA library, ensuring sufficient coverage for robust demultiplexing and transcript quantification.

**Bioinformatic and statistical analysis**

FASTQ files were processed with the BD Rhapsody™ WTA Analysis Pipeline (Seven Bridges, Revision 8), using an adapted VDJ alignment module to retrieve LMO sequences. This produced filtered and unfiltered expression matrices, BAM files, and quality-control reports. Unique molecular identifier (UMI) matrices were then analyzed in R (v4.4.0) using the Seurat (v5.1.0) and MULTI-seq packages. Each cell was required to contain ≥100 LMO counts and to express at least two-thirds of a specific SBC relative to total LMO counts. For transcriptome data, high-quality cells were defined as those with 1,000–10,000 detected genes and <0.4% mitochondrial reads.

Normalized datasets were subjected to dimensionality reduction via principal component analysis (PCA), and significant PCs were selected based on the inflection point of the elbow plot. Unsupervised Louvain clustering and UMAP visualization were performed with 20 PCs and 50 neighbors. For integrated analyses, the Seurat integration workflow was applied to combine samples across conditions. Marker genes for each cluster were identified using ROC tests, and classification power was assessed by the area under the ROC curve. Identified markers were cross-referenced with known developmental regulators (e.g., Guillot et al., 2021) to validate cell-type assignments.

### ADDITIONNAL RESOURCES

### QUANTIFICATION AND STATISTICAL ANALYSIS

### KEY RESOURCES TABLE

#### TABLE FOR AUTHOR TO COMPLETE

***Please do not add custom subheadings.*** *If you wish to make an entry that does not fall into one of the subheadings below, please contact your handling editor or add it under the “other*” subheading*.* ***Any subheadings not relevant to your study can be skipped.*** *(****NOTE:*** *references should be in numbered style, e.g., Smith et al.^1^)*

**Key resources table**

| **REAGENT or RESOURCE** | **SOURCE** | **IDENTIFIER** |
| --- | --- | --- |
| **Antibodies** | | |
| **Bacterial and virus strains** | | |
| **Biological samples** |  |  |
| **Chemicals, peptides, and recombinant proteins** | | |
| FVS 660 | BD | #564405 |
| MULTI-Seq LMO Lig-Anchor | Gene Link | #26-2202-02 |
| MULTI-Seq LMO Palm Co-Anchor | Gene Link | #26-2204-02 |
| Accutase | Gibco | #A1110501 |
| **Critical commercial assays** | | |
| BD Rhapsody^TM^ Cartridge Reagent Kit | BD | #633731 |
| BD Rhapsody^TM^ Cartridge Kit | BD | #633733 |
| BD Rhapsody^TM^ cDNA kit | BD | #633773 |
| BD Rhapsody^TM^ WTA Amplification Kit | BD | #633801 |
| AgencourtR AMPureR XP magnetic beads | Beckman Coulter Life Science | #A63880 |
| **Deposited data** | | |
| GSE |  |  |
| **Experimental models: Cell lines** | | |
| **Experimental models: Organisms/strains** | | |
| Gallus gallus / JA 57 strain | EARL Les Bruyères, Dangers, France |  |
| **Oligonucleotides** | | |
| See supplementary Table1 |  |  |
| **Recombinant DNA** | | |
| **Software and algorithms** | | |
| BD Rhapsody™ WTA Analysis Pipeline run (Revision 8) | Seven Bridges | Sevenbridges.com |
| Rstudio | R 4.4.0 | https://posit.co/download/rstudio-desktop/ |
| Seurat | Version 5.1.0 | https://satijalab.org/seurat/ |
| **Other** | | |

***PHYSICAL SCIENCES***

| **REAGENT or RESOURCE** | **SOURCE** | **IDENTIFIER** |
| --- | --- | --- |
| Chemicals, peptides, and recombinant proteins | | |
| QD605 streptavidin conjugated quantum dot | Thermo Fisher Scientific | Cat#Q10101MP |
| Platinum black | Sigma-Aldrich | Cat#205915 |
| Sodium formate BioUltra, ≥99.0% (NT) | Sigma-Aldrich | Cat#71359 |
| Chloramphenicol | Sigma-Aldrich | Cat#C0378 |
| Carbon dioxide (^13^C, 99%) (<2% ^18^O) | Cambridge Isotope Laboratories | CLM-185-5 |
| Poly(vinylidene fluoride-co-hexafluoropropylene) | Sigma-Aldrich | 427179 |
| PTFE Hydrophilic Membrane Filters, 0.22 mm, 90 mm | Scientificfilters.com/Tisch Scientific | SF13842 |
| Critical commercial assays | | |
| Folic Acid (FA) ELISA kit | Alpha Diagnostic International | Cat# 0365-0B9 |
| TMT10plex Isobaric Label Reagent Set | Thermo Fisher | A37725 |
| Surface Plasmon Resonance CM5 kit | GE Healthcare | Cat#29104988 |
| NanoBRET Target Engagement K-5 kit | Promega | Cat#N2500 |
| Deposited data | | |
| B-RAF RBD (apo) structure | This paper | PDB: 5J17 |
| Structure of compound 5 | This paper; Cambridge Crystallographic Data Center | CCDC: 2016466 |
| Code for constraints-based modeling and analysis of autotrophic *E. coli* | This paper | https://gitlab.com/elad.noor/sloppy/tree/master/rubisco |
| Software and algorithms | | |
| Gaussian09 | Frish et al.^1^ | https://gaussian.com |
| Python version 2.7 | Python Software Foundation | [https://www.python.org](https://www.python.org/) |
| ChemDraw Professional 18.0 | PerkinElmer | <https://www.perkinelmer.com/category/chemdraw> |
| Weighted Maximal Information Component Analysis v0.9 | Rau et al.^2^ | https://github.com/ChristophRau/wMICA |
| Other | | |
| DASGIP MX4/4 Gas Mixing Module for 4 Vessels with a Mass Flow Controller | Eppendorf | Cat#76DGMX44 |
| Agilent 1200 series HPLC | Agilent Technologies | https://www.agilent.com/en/products/liquid-chromatography |
| PHI Quantera II XPS | ULVAC-PHI, Inc. | https://www.ulvac-phi.com/en/products/xps/phi-quantera-ii/ |
