## Supplementary Tables for "Enhanced Multiplexed Single-Cell RNA-Sequencing for Accurate Detection of Treatment Effects Without Batch Correction in the Avian Embryonic Model"

### Supplementary Table 1: Design of LMO labeling testing of chick cells

| | TEST 1 200nM | TEST 2 20nM | TEST 3 2nM |
|---|---|---|---|
| Cell number | 500K | 500K | 500K |
| Final volume of cells incubation (µL) | 200µL | 200µL | 200µL |
| Final concentration LMOs (nM) | 200nM | 20nM | 2nM |
| LMOs (µL) + <br><br> Probe Probe anti LMOs Atto550 (µL) + <br><br> FVS 640 nm (µL) <br><br> ([LMOs]i = 2µM) | - 20µL LMO Anchor + probe anti LMO atto550 qsp 159µL PBS 1X ;<br>- add on washed dissociated cells<br>- 5min on ice ;<br>- add 20µL LMO CoAnchor.<br>- gentle mixing.<br>- 5min on ice ;<br>- Centrifugation 500xg 10 min, take off supernatant<br>- then add 200µL HBSS (FACS melody Analysis) | - | - |
| LMOs (µL) + <br><br> Probe Probe anti LMOs Atto550 (µL) + <br><br> FVS 660 nm (µL) <br><br> ([LMOs]i = 0,2µM) | - | - 20µL LMO Anchor + probe anti LMO atto550 qsp 159µL PBS 1X ;<br>- add on washed dissociated cells<br>- 5min on ice ;<br>- add 20µL LMO CoAnchor.<br>- gentle mixing.<br>- 5min on ice ;<br>- Centrifugation 500xg 10 min, take off supernatant<br>- then add 200µL HBSS (FACS melody Analysis) | - |

| | | | |
|---|---|---|---|
| LMOs (µL) + <br><br> Probe Probe anti LMOs Atto550 (µL) + <br><br> FVS 660 nm (µL) <br><br> ([LMOs]i = 0,2µM) | - | - | - 2µL LMO Anchor + probe anti LMO atto550 qsp 159µL PBS 1X ; <br> - add on washed dissociated cells <br> - 5min on ice ; <br> - add 2µL LMO CoAnchor. <br> - gentle mixing. <br> - 5min on ice ; <br> - Centrifugation 500xg 10 min, take off supernatant <br> - then add 200µL HBSS <br> (FACS melody Analysis) |

### Supplementary Table 2: Sequences of Sample Barcodes (SBC)

| Name | Barcode Sequen | Oligo sequence (5' -> 3') |
|---|---|---|
| **SBC1** | CCACAATG | CCTTGGCACCCGAGAATTCCACCACAATGAAAAAAAAAAAA AAAAAAAAAAAAAAAAAA |
| **SBC3** | TGAGACCT | CCTTGGCACCCGAGAATTCCATGAGACCTAAAAAAAAAAAA AAAAAAAAAAAAAAAAAA |
| **SBC7** | GAAAAGGG | CCTTGGCACCCGAGAATTCCAGAAAAGGGAAAAAAAAAAAA AAAAAAAAAAAAAAAAAA |

### Supplementary Table 3: Sequences of specific primers used for library amplification

| Name | Sequence (5' -> 3') |
|---|---|
| Custom-LibraryP7-R1 | CAAGCAGAAGACGGCATACGAGATAGCGTAGCGTGACTGGAGTTCAGACGTGTGCTCTTCCGATCTCCT TGGCACCCGAGAATTCCA |
| Custom-LibraryP7-R2 | CAAGCAGAAGACGGCATACGAGATCAGCCTCGGTGACTGGAGTTCAGACGTGTGCTCTTCCGATCTCCT TGGCACCCGAGAATTCCA |
| Custom-LibraryP7-R3 | CAAGCAGAAGACGGCATACGAGATCGAGTAATGTGACTGGAGTTCAGACGTGTGCTCTTCCGATCTCCT TGGCACCCGAGAATTCCA |
| Custom-LibraryP7-R4 | CAAGCAGAAGACGGCATACGAGATTCTCCGGAGTGACTGGAGTTCAGACGTGTGCTCTTCCGATCTCCT TGGCACCCGAGAATTCCA |

| | |
|---|---|
| Multiseq-additive-primerandProbe | CCTTGGCACCCGAGAATTCC |
